# A comparative study of statistical methods for identifying differentially expressed genes in spatial transcriptomics

**DOI:** 10.1101/2025.02.17.638726

**Authors:** Yishan Wang, Chenxuan Zang, Ziyi Li, Charles C. Guo, Dejian Lai, Peng Wei

## Abstract

Spatial transcriptomics (ST) provides unprecedented insights into gene expression patterns while retaining spatial context, making it a valuable tool for understanding complex tissue architectures, such as those found in cancers. Seurat, by far the most popular tool for analyzing ST data, uses the Wilcoxon rank-sum test by default for differential expression analysis. However, as a nonparametric method that disregards spatial correlations, the Wilcoxon test can lead to inflated false positive rates and misleading findings. This limitation highlights the need for a more robust statistical approach that effectively incorporates spatial correlations. To this end, we propose a Generalized Estimating Equations (GEE) framework as a robust solution for differential gene expression analysis in ST. We conducted a comprehensive comparison of the GEE-based tests with existing methods, including the Wilcoxon rank-sum test and z-test. By appropriately accounting for spatial correlations, extensive simulations showed that the GEE test with robust standard error, referred to as the Independent GEE, demonstrated superior Type I error control and comparable power relative to other methods. Applications to ST datasets from breast and prostate cancer showed poor calibration of the p-values and potential false positive findings from the Wilcoxon rank-sum test. Our comparative study based on simulations and real data applications suggests that the Independent GEE test is well-suited for ST data, offering more accurate identification of biologically relevant gene expression changes and complementing the Wilcoxon rank-sum test. We have implemented the proposed method in R package “SpatialGEE”, available on GitHub.

**Author Summary:** Spatial transcriptomics (ST) provides unprecedented insights into gene expression patterns while retaining spatial context, making it a valuable tool for studying complex tissue architectures and disease etiology. Seurat, a widely used software tool for analyzing ST data, relies on the Wilcoxon rank-sum test for differential expression analysis. However, this test ignores spatial correlations, leading to inaccurate control of false positive rates and misleading findings. This limitation highlights the need for a more robust statistical approach that effectively incorporates spatial correlations. To this end, we have proposed a Generalized Estimating Equation (GEE) framework as a robust solution for differential gene expression analysis in ST. By appropriately accounting for spatial correlations, extensive simulations showed that the GEE-based test demonstrated superior false positive rate control and comparable power relative to other methods. Applications to ST datasets from breast and prostate cancer showed potential false positive findings from the Wilcoxon rank-sum test. We recommend the GEE method to be a useful complement to the Wilcoxon rank-sum test. We have implemented the proposed method in R package “SpatialGEE”, available on GitHub.

## 1. Introduction

Spatial transcriptomics (ST) is an emerging high-throughput technology that enables genome-wide profiling of gene expression while preserving the spatial context within tissues, providing critical insights into cellular function, tissue organization, and intercellular communication (Ståhl et al., 2016; Svensson et al., 2018; Erickson et al., 2022; Tian et al., 2023; Jiang et al., 2024; Ma & Zhou, 2024; Shah et al., 2024). A fundamental analytical goal in ST studies is to identify differentially expressed (DE) genes across pathological regions or disease grades, for example, between ductal carcinoma in situ and invasive carcinoma in breast cancer, to elucidate spatially resolved molecular mechanisms underlying development and disease.

Several statistical frameworks have been developed for ST analysis, each addressing distinct aspects of spatial expression. SpatialDE (Svensson et al., 2018) and related methods such as SPARK (Sun et al., 2020) aim to identify spatially variable genes whose expression levels vary smoothly across the entire slide/tissue coordinates. C-SIDE (Cable et al., 2022) extends this concept to cell type–specific differential expression across regions by integrating ST data with single-cell RNA-seq references to infer cell type compositions within each spot, followed by in silico differential testing on the deconvolved expression profiles. In contrast, studies focused on comparing predefined histological or pathological regions typically rely on classical nonparametric tests such as the Wilcoxon rank-sum test (Wilcoxon, 1945), which remains a default choice due to its computational simplicity and broad availability in analysis pipelines such as Seurat (Butler et al., 2018; Stuart et al., 2019; Hao et al., 2024).

More recently, spatialGE (Ospina et al., 2022, 2025) has expanded this toolkit by integrating statistical and spatial modeling approaches for DE analysis in ST data. spatialGE first performs non-spatial tests (e.g., Wilcoxon rank-sum or two-sample t-test) to identify candidate DE genes and subsequently applies a spatial linear mixed model (LMM) with an exponential covariance structure only to this subset of genes. However, ST data are typically zero-inflated counts, for which t-tests and LMMs are not well-suited, as both rely on normality assumptions that are violated by count-based or sparsely expressed genes. In this context, we also considered the two-sample z-test as a more general alternative to the t-test. Unlike the t-test, the z-test does not assume normality and is applicable to arbitrarily distributed data, including counts or binary observations, as long as the sample size in each comparison group is sufficiently large (e.g., >50 spots per pathological region, which is typical in ST data). Nevertheless, both the t- and z-tests, as well as the Wilcoxon rank-sum test, assume independent observations; when spatial correlations are present, violations of this assumption can inflate or deflate the Type I error rate, depending on the magnitude and structure of pairwise correlations among observations (Lehmann and Romano, 2022).

Despite these methodological advances, there has been no systematic evaluation of existing approaches for DE analysis in ST data with respect to Type I error control, power, numerical stability, and computational efficiency, highlighting an urgent need for comprehensive benchmarking and methodological innovation in this rapidly evolving field.

Our preliminary analyses of real datasets and simulated data have revealed limitations of the Wilcoxon rank-sum test when applied to the spatially-correlated ST data. Specifically, we observed that the Wilcoxon rank-sum test tends to have inflated Type I error rates in the presence of strong spatial correlations, resulting in an increased number of false positives. This issue raises concerns about the validity of power estimates and compromises the credibility of findings derived from spatial transcriptomic studies. Given that most transcriptomic data exhibit inherent spatial dependencies, Wilcoxon rank-sum test’s assumption of independence between observations is frequently violated, emphasizing the need for alternative approaches that are more suitable for spatially structured data.

In this study, we considered two potential approaches: generalized linear mixed model (GLMM) and generalized estimating equations (GEE). The GLMM has been considered by many investigators as the gold standard for analysis of correlated data, since they are flexible in modeling complex dependency structures, thus allowing to account for both fixed and random effects (Breslow & Clayton, 1993). However, the GLMM can be computationally challenging in high-dimensional ST due to the large number of parameters involved and zero-inflated count data, which often leads to convergence issues and significant computational demands. To mitigate these computational challenges, we also investigate the GEE, a marginal modeling framework that offers a balance of robustness with computational efficiency (Liang & Zeger, 1986; Zeger et al., 1988). Unlike the GLMM, which models random effects explicitly, the GEEs use a “working” correlation matrix to effectively account for the spatial dependence between observations. More precisely, we first proposed and implemented the generalized score test (GST) (Rao, 1948; Liang & Zeger, 1986; Zhang et al., 2014) within the GEE framework. We then compared two variations of the GEEs: the commonly used robust Wald test (White, 1980; Huber, 1967) and the GST. The robust Wald test requires fitting the model under the alternative hypothesis and uses a robust sandwich estimator for the standard error to account for correlations within the data. In contrast, the GST requires only the fitting of the null model, which enhances numerical stability.

We excluded the GLMM in our comparison due to its computational intensity and convergence issues when applied to zero-inflated ST data. In addition to simulation studies focusing on zero-inflated sparse count data, we applied the Wilcoxon rank-sum test, two-sample z-test, and a family of GEE-based tests to real datasets from breast and prostate cancer from 10x Genomics (Figure 1a and Figure 1b) to evaluate their effectiveness in detecting DE genes between tumor and normal tissues. The quantile-quantile (QQ) plots revealed poor calibration of p-values by the Wilcoxon rank-sum test, underscoring the need for a comprehensive comparative study in a statistically principled framework.

**Figure 1.**
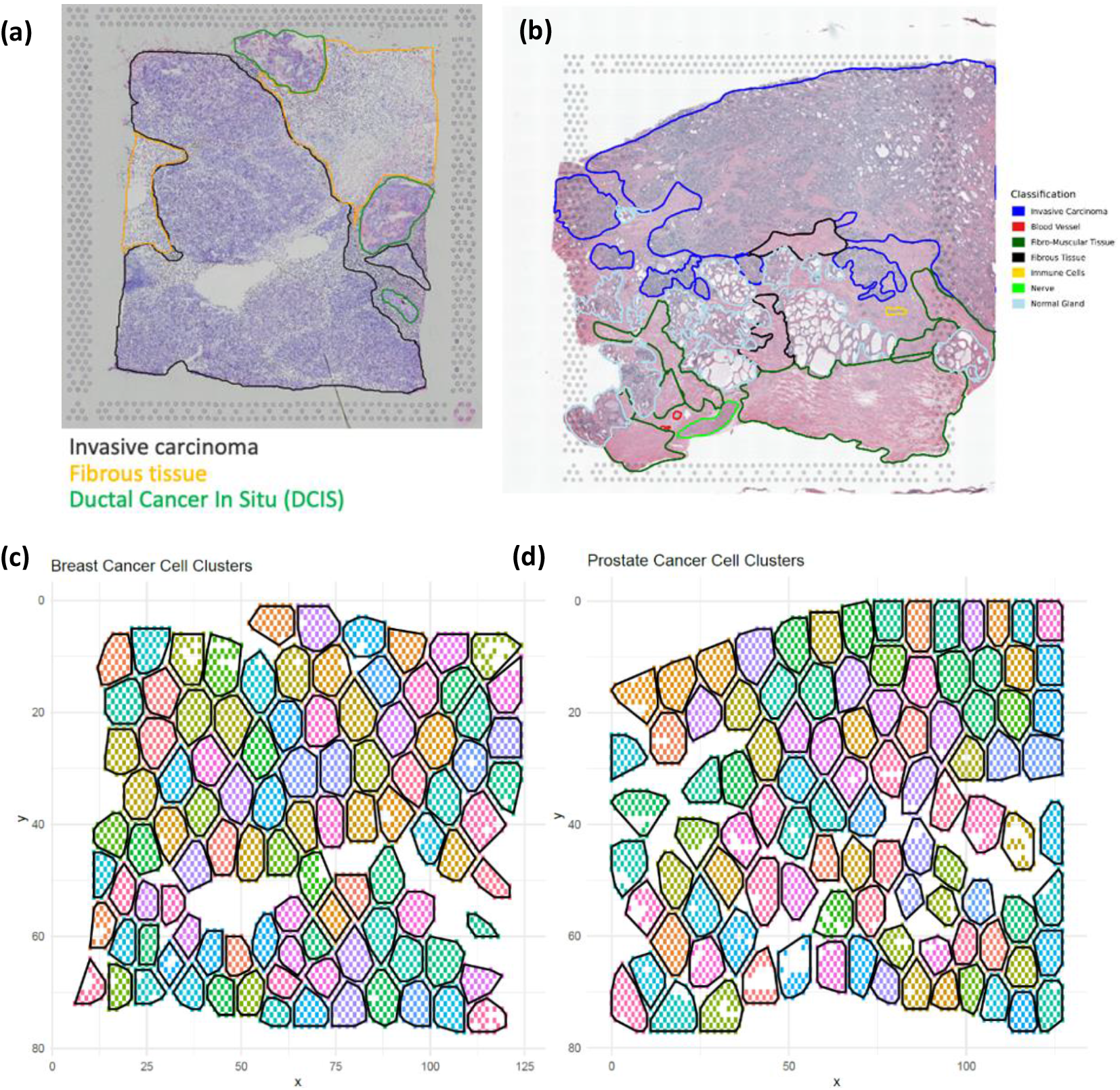
H&E-stained images and spatial clusters for breast and prostate cancer. **(a)** H&E-stained image with pathology labels for breast cancer (10x Genomics). **(b)** H&E-stained image with pathology labels for prostate cancer (10x Genomics; Zang et al., 2024). **(c)** Spatial clusters (n = 100) for breast cancer. **(d)** Spatial clusters (n = 100) for prostate cancer.

The rest of the paper is organized as follows. Section 2 reviews and introduces statistical methods under comparison for detecting DE genes in ST data, including Wilcoxon rank-sum test, GLMM, and robust Wald test and GST in the GEE framework. Section 3 presents a comprehensive simulation study to compare Type I error rates and power, followed by applications to a breast cancer and a prostate cancer ST dataset. We conclude with a discussion of our main findings and future directions.

## 2. Materials and Methods

### 2.1. Wilcoxon Rank-Sum Test

We begin with the Wilcoxon rank-sum test due to its use as the default method in popular ST data analysis software suites like Seurat and SpatialGE. Its simplicity and computational efficiency make it the most common method for detecting DE genes across pathological grades in ST data.

Consider a tissue containing two pathology grades: Grade A and Grade B. Each spatial location *i* (*i* = 1, …, *n*) is represented by a spot that includes multiple cells, along with associated gene expression data. Let *X* and *Y* represent the expression levels of a particular gene in Grades A and B, respectively. The number of spatial locations in Grade A is denoted by *n*_*X*_, and the number of spatial locations in Grade B is denoted by *n*_*Y*_.

The Wilcoxon rank-sum test assesses differential expression between the two grades by ranking all observations, calculating the sum of ranks for each grade, and computing the test statistic based on these sums. The rank of the *i*th observation when combining and ordering all observations from both grades is represented by *R*_*i*_. The sum of ranks for Grade A is *W*_*X*_ = ∑ *R*_*i,X*_, where *R*_*i,X*_ is the rank of the *i*th observation in Grade A. Similarly, the sum of ranks for Grade B is *W*_*Y*_ = ∑ *R*_*i,Y*_.

To evaluate the difference between the distributions of *X* and *Y*, the test statistic *W* = *min*(*W*_*X*_, *W*_*Y*_) is used. When the sample sizes (*n*_*X*_ and *n*_*Y*_) are sufficiently large, the distribution of *W* under the null hypothesis *H*_0_: the distributions of *X* and *Y*are identical, can be approximated by a normal distribution with mean 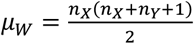 and variance 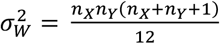 .The standardized test statistic is then calculated as 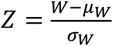, and the two-sided p-value is obtained as *p* − *value* = 2 × (1 − *Φ*(|*Z*|)), where Φ(|*Z*|) represents the cumulative distribution function (CDF) of the standard normal distribution for the absolute value of *Z*.

Despite its common use in practice, the Wilcoxon test assumes independence between observations, an assumption often violated in spatially correlated data. This can lead to either inflated or deflated Type I error rates depending on the correlation patterns (Datta and Satten, 2005), highlighting the need to explore alternative methods that account for spatial dependencies. To control the false discovery rate (FDR), the Benjamini-Hochberg (BH) procedure can be applied for multiple testing in genome-wide scans (Benjamini & Hochberg, 1995); to control the family-wise error rate (FWER), we apply the Bonferroni procedure.

### 2.2. Introduction to GLMM and GEE for Clustered Count Data

In spatial transcriptomics, gene expression measurements often exhibit inherent correlation structures: spots within the same tissue region or spot tend to share biological and technical variability. Neglecting such correlations can lead to biased parameter estimates, underestimated standard errors, and inflated Type I error rates. To address this issue, two major statistical frameworks have been widely employed for analyzing correlated and clustered outcomes: the GLMM and the GEE (McCulloch et al., 2008; Liang and Zeger, 1986).

The GLMM extends the generalized linear model (GLM) by incorporating random effects to capture cluster- or subject-specific variability while preserving the link between the response mean and covariates through a canonical link function. GLMMs assume that the conditional distribution of the response belongs to the exponential family (e.g., Poisson for count data) and specify a distribution for the random effects, commonly Gaussian. This likelihood-based framework allows for flexible modeling of within-cluster correlations and enables subject-level inference. However, when applied to large-scale and zero-inflated count data, such as those commonly observed in spatial transcriptomics, GLMMs frequently encounter computational instability and convergence failures, particularly when estimating weak spatial correlations within a high-dimensional likelihood framework (Bolker et al., 2009; Breslow and Clayton, 1993).

In contrast, the GEE adopts a semi-parametric, population-averaged approach by specifying only the mean model and a “working” correlation structure that approximates within-cluster dependence (Liang and Zeger, 1986). Rather than maximizing a likelihood, GEE estimates regression parameters by solving a set of quasi-score equations, and employs a robust (sandwich) variance estimator to maintain consistency, i.e., the estimation bias approaches 0 as the number of clusters grow, even when the working correlation is misspecified. This robustness and computational efficiency make GEE particularly suitable for large, correlated, and high-dimensional count data, where fully parametric models may be computationally prohibitive or numerically unstable.

Overall, GLMM and GEE offer complementary advantages for modeling clustered count data: GLMM provides a fully specified hierarchical likelihood that enables cluster-specific inference but may suffer from scalability issues, while GEE emphasizes robust, population-level inference with reduced computational burden. We next introduce the GLMM and GEE in the context of modeling spatially correlated ST count data.

### 2.3. GLMM for spatially correlated ST count data

We explored the GLMM here because it is considered the “gold standard” for analyzing spatially correlated data. Its flexibility in modeling fixed and random effects allows it to account for spatial dependencies, making it initially promising for identifying DE genes across pathology grades.

Let *Y*_*ij*_ (*i* = 1, …, *n*; *j* = 1, …, *p*) represent the gene expression count for gene *j* at spatial location *i*. Define *X*_*i*_ as a binary dummy variable for pathology grade at location *i* (0 for Grade A, 1 for Grade B). The spatial coordinates are denoted by *s*_*i*_ = (*s*_*i*1_, *s*_*i*2_) for spatial location *i*.

Following the SPARK (Sun et al, 2020) formulation, we model the observed count data *Y*_*ij*_ via the following GLMM for the Poisson family:

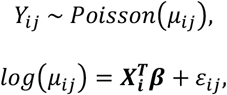

where *μ*_*ij*_ is the expected count for gene *j* at location *i* and; 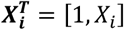 is the design matrix for fixed effects; ***β*** = [*β*_*j*0_, *β*_*j*1_]^*T*^ is the vector of fixed effect coefficients, where *β*_*j*0_ is the intercept and *β*_*j*1_ is the effect of Grade B compared to Grade A. The random effect *ε*_*ij*_ is assumed to follow a normal distribution, 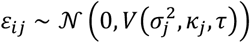, representing the random effect for gene *j* at the *i*th spatial location.

For a given gene *j*, the spatial covariance matrix 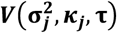 is defined based on the distances between pairs of spatial locations, and 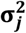 represents a vector of variance components (Li et al., 2009). The (*i, i*^′^)th element of 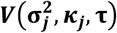 is given by 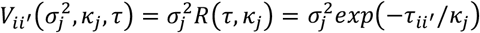, where *τ*_*ii*_ ′ = ||*s*_*i*_ − *s*_*i*_ ′ || denotes the Euclidean distance between two spatial locations *i* and *i*^′^. Here, *κ*_*j*_ > 0 is a parameter that determines the rate of decay in correlation with distance, with larger values of *κ*_*j*_ indicating stronger correlations and smaller semi-variances (1 − *exp*(−*τ*/*κ*)). The exponential spatial structure used here is a specific case of the Matérn correlation structure 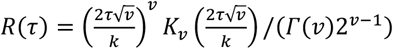 when *ν* = 0.5.

To test for DE genes across the two pathology grades, we test the null hypothesis *H*_0_: *β*_*j*1_ = 0 against the alternative hypothesis *H*_*a*_: *β*_*j*1_ ≠ 0. This tests whether the expected count of gene expression significantly differs between the pathology grades after accounting for spatial correlation. Statistical inference is made using likelihood ratio tests based on the maximum likelihood estimation of the GLMM. For example, to test *H*_0_: *β*_*j*1_ = 0, i.e., gene *j* is not differentially expressed across Grade A and Grade B, we compare the full model with the null model *log*(*μ*_*ij*_) = *β*_0_ + *ε*_*ij*_. DE genes are then identified by applying the Bonferroni procedure to control the FWER.

### 2.4. GEE for spatially correlated ST count data

Instead of explicitly modeling the spatial correlation structure using random effects, the GEE model uses a “working” correlation matrix to account for the spatial dependence between observations. We adopted the GEE model with an independent working correlation structure by dividing the whole ST tissue into *m* spatial clusters (Figure 1c and Figure 1d) using K-means clustering (MacQueen, 1967). Specifically, we used spatial coordinates (x, y) as the inputs to the K-means algorithm. The purpose of this clustering step is not to identify biological subtypes, but rather to define spatially contiguous groups of neighboring spots that serve as working “clusters” in the GEE framework. This procedure is conceptually similar to dividing the tissue into local spatial neighborhoods, but the K-means algorithm provides flexibility in capturing irregular tissue shapes and densities, in contrast to a fixed grid partitioning. The mean of *Y*_*ij*_, denoted by *μ*_*ij*_, is linked to the covariates through a log link function, and the model is specified as:

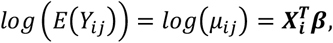

where 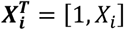 is the design matrix; ***β*** = [*β*_*j*0_, *β*_*j*1_] ^*T*^ is the vector of fixed effect coefficients.

The parameters ***β*** are estimated by solving the GEE:

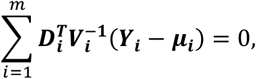

where ***D***_***i***_ is the derivative of the mean response with respect to ***β*** in the cluster *i* 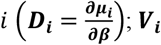 is the variance-covariance matrix of responses in the cluster *i*, which is a function of the working correlation matrix 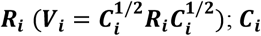 is a diagonal matrix that includes the variances of the individual observations within the cluster *i*; ***Y***_***i***_ is the response vector in cluster *i*; ***μ***_***i***_ is the mean vector in the cluster *i*. The robust variance estimates are used to make inferences for the estimated coefficients from the GEE. The robust variance estimate for the estimated coefficients is calculated by the “sandwich” estimator:

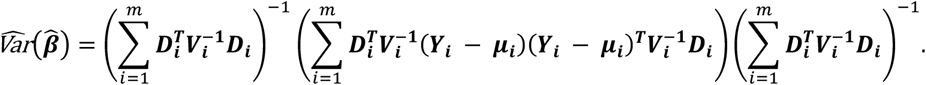

To provide a more efficient model fitting, the GEE uses an independent working correlation matrix ***R*** (an *n* × *n* matrix where *n* is the number of spatial locations). The advantage of the GEE framework is that it produces consistent and asymptotically normal estimates of the parameters even if the working correlation structure is incorrectly specified (Liang & Zeger, 1986).

#### 2.4.1. Robust Wald Test

The robust Wald test requires fitting the full GEE model under the alternative hypotheses to identify DE genes across the two pathology grades. This approach is the default test implemented in all currently available R packages associated with the GEE framework. To test *H*_0_: *β*_*j*1_ = 0 against *H*_*a*_: *β*_*j*1_ ≠ 0, the robust Wald test statistic *W* is computed as:

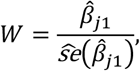

where 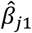 is the estimated coefficient and 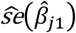 is the robust standard error, which is the square root of the “sandwich” variance estimator. *W* approximately follows a standard normal distribution under the null hypothesis when the number of clusters *m* is large.

A special case of the GEE with the robust Wald test is to treat each spatial location as a unique cluster, i.e., cluster number *m* = *n* assuming independence among all spatial locations. This model is referred to as the Independent GEE. Although it is conceptually similar to a two-sample z-test due to the working assumption of independent observations, it uses the sandwich estimator-based robust standard error to effectively correct for the working independence, yet often wrong, assumption. As to be shown in the simulation study, the Independent GEE controls the Type I error rate precisely despite the spatially correlated observations, while the two-sample z-test has inflated Type I error rate.

#### 2.4.2. Generalized Score Test (GST)

We further propose the GST as an alternative to the robust Wald test, sharing the same asymptotic properties but requiring only model fitting under the null hypothesis, making it more numerically stable. The asymptomatic equivalence between the GST statistic and the robust Wald statistic holds only as the number of spatial clusters is large (or approaches infinity in theory). However, with a finite number of clusters, as observed in real ST data, these tests can yield different Type I error and power results (Boos & Stefanski, 2013), supporting the need for more numerically robust methods in practical settings.

The score function 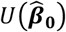 is the derivative of the quasi-loglikelihood with respect to the parameters ***β***, evaluated at the estimated parameters under the null hypothesis. Specifically, the score function 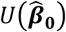 is given by:

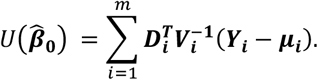

The robust variance estimates of the score statistic, 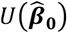, is also estimated using the “sandwich” estimator 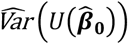 . To test *H*_0_: *β*_*j*1_ = 0 against *H*_*a*_: *β*_*j*1_ ≠ 0, the GST statistic *S* is computed as:

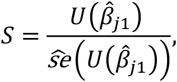

where 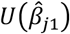 is the score function evaluated at the estimated parameter 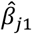 under the null hypothesis, and 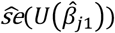 is the robust standard error of the score function. The test statistic *S* approximately follows a standard normal distribution under the null hypothesis when the number of clusters *m* is large.

## 3. Results

### 3.1. Simulation Studies

We conducted extensive simulation studies that closely resembled the conditions found in real data, including spatial structure and zero-inflated characteristics, for rigorously comparing the performance of different statistical methods in the context of ST. First, we fitted the GLMM described in Section 2.3 to a breast cancer ST dataset from 10x Genomics, to be detailed in Section 3.2.1. This gave an estimation of the two key spatial parameters: *σ*^2^ and *κ*, where *σ*^2^ is the spatial variance that captures variability in gene expression across different spatial locations and κ is a decay parameter that controls the rate of decay in correlation with distance between spatial locations. These estimates were calculated from the fitted results of the GLMM for genes that had converged. The estimated parameters were subsequently used to generate simulated data that preserved the realistic spatial structure found in biological samples, closely resembling real ST datasets. Specifically, we employed the same GLMM structure by applying the estimated *σ*^2^ and *κ* to ensure spatial correlation, while introducing zero-inflation through a small intercept (*β*_0_) in the model. This approach preserved the spatial heterogeneity observed in real ST data and captured the intrinsic sparsity characteristic of biological samples. To better capture the range of spatial variability present in real tissues, we took set of three levels: 25th percentile, median, and 95th percentile in variation of the spatial parameters, thus representing the spectrum of variance in spatial gene expression and spatial correlation strengths from weak to strong. Regarding this, we considered three scenarios: (1) the 25th percentile of *σ*^2^ and the 25th percentile of *κ*, (2) the median of *σ*^2^ and the median of *κ*, and (3) the 95th percentile of *σ*^2^ and the 95th percentile of *κ*.

These simulations target two aspects of model performance: Type I error rate control and statistical power. Type I error rate control is important to assess the reliability of the results, especially in ensuring that false positives are controlled at the nominal level. For each statistical method, we produced 100,000 replicates to evaluate how well the different approaches could control the rate of false positives at different levels of significance. The cut-offs ranged from α = 0.01 to the more stringent α = 0.0001. It is in genome-wide analyses that such stringent levels of α are required, due to the large number of tests being conducted simultaneously. On the other hand, owing to the computational intensive nature of these simulations, we did not explore even lower levels of α.

In addition to Type I error control, we investigated the statistical power of each method, which is indicative of the number of DE genes correctly identified. We generated 1,000 replicates to compare the power of different statistical approaches in the detection of true signals while ensuring Type I error rates remain acceptable. Because GEE rely on asymptotic behavior and thus require large numbers of clusters to make proper inferences, under each scenario we set the number of spatial clusters to either 25 or 100. More precisely, we investigated the impact of the number of spatial clusters on the performance of the GEE. When the model was configured to treat each spatial location as an independent cluster, it reduced to a special case we termed Independent GEE. Despite its conceptual similarity to a two-sample z-test due to the working assumption of independent observations, the Independent GEE uses the sandwich estimator-based robust standard error to effectively correct for the working independence, yet often wrong, assumption.

Of note, although theoretically relevant for modeling correlated data, we excluded the GLMM from the final simulations and comparisons. We made this decision due to its inability to converge for many genes (about 43%) and extremely demanding computational runtime (some genes took more than 10 hours to finish model fitting) when applied to the breast cancer ST data to be detailed in Section 3.2.1. The zero-inflated count nature of the real data resulted in weak spatial correlations, leading to convergence issues for the GLMM when estimating spatial parameters in the full likelihood framework. Failure to converge and yield stable results from the GLMM highlighted its limitations in handling the computational and numerical challenges in the context of ST data. Consequently, our comparisons focused on five methods: the Wilcoxon rank-sum test, two-sample z-test, the GEE with the robust Wald test, the GEE with GST, and the Independent GEE.

The simulation results are presented in Table 1 for Type I error rate comparison and in Figure 2 for power comparison. We found that when the number of spatial clusters was sufficiently large – *m* =100 clusters in our case, the GEE with robust Wald test and GEE with GST in general improved Type I error control compared to *m* = 25; nevertheless, both tests showed either deflated or inflated Type I error rates across spatial correlation scenarios even with *m* =100. On the other hand, Independent GEE with the robust Wald test showed consistently satisfactory control of the Type I error rates across spatial correlation scenarios, even under the strong spatial correlations where all other tests failed to control the Type I error rate satisfactorily. The Wilcoxon rank-sum test had the empirical Type I error rates close to the nominal levels under weak and moderate spatial correlation scenarios but had inflated Type I error rates across α levels when the spatial correlation was strong. Finally, the two-sample z-test could control the Type I error rates only under the weak spatial correlation scenario and had severely inflated Type I error rates under moderate and strong spatial correlations. Due to its inflated Type I error rate, we excluded the two-sample z-test in the power comparison. Figure 2 shows the power comparison under α = 0.001. All tests performed similarly except for when the number of clusters *m* = 25 the GEE with GST appeared to be the most conservative test with the lowest statistical power (Figure 2 a, c, and d). When *m* = 100, it became on par with the other tests (Figure 2 b, d, and e). In summary, our simulation studies suggest that the Independent GEE is the most reliable method for detecting DE genes in ST data, serving as a valuable complement to the Wilcoxon rank-sum test.

**Table 1.** Type I error comparison of GEE with robust Wald test, GEE with GST, Independent GEE with robust Wald test, Wilcoxon rank-sum test, and z-test across different spatial scenarios and spatial cluster numbers. Type I error was evaluated at significance levels of α = 0.01, 0.001, and 0.0001 across 100,000 simulation replications. Inflated or deflated empirical Type I error rates (larger or smaller than the nominal α by 50% or more) are in boldface.

| | | Significance level $\alpha$ | | |
| --- | --- | --- | --- | --- |
| # of Clusters | Tests | 0.01 | 0.001 | 0.0001 |
| Weak Spatial Correlation and Spatial Variance (Scenario 1) |  |  |  |  |
| 25 | GEE with GST | 0.008 | **0.0003** | **0.00001** |
| 25 | GEE with Wald test | **0.022** | **0.0049** | **0.00138** |
| 100 | GEE with GST | 0.010 | 0.0010 | 0.00013 |
| 100 | GEE with Wald test | 0.011 | 0.0014 | **0.00026** |
| - | Independent GEE with Wald test | 0.008 | 0.0008 | 0.00009 |
| - | Wilcoxon rank-sum test | 0.008 | 0.0009 | 0.00008 |
| - | z-test | 0.009 | 0.0011 | 0.00012 |
| Moderate Spatial Correlation and Spatial Variance (Scenario 2) |  |  |  |  |
| 25 | GEE with GST | 0.010 | **0.0002** | **0** |
| 25 | GEE with Wald test | **0.025** | **0.0057** | **0.00166** |
| 100 | GEE with GST | **0.019** | **0.0023** | **0.00028** |
| 100 | GEE with Wald test | **0.017** | **0.0023** | **0.00039** |
| - | Independent GEE with Wald test | 0.013 | 0.0011 | 0.00009 |
| - | Wilcoxon rank-sum test | 0.013 | 0.0012 | 0.00008 |
| - | z-test | **0.020** | **0.0030** | **0.00041** |
| Strong Spatial Correlation and Spatial Variance (Scenario 3) |  |  |  |  |
| 25 | GEE with GST | **0.00001** | **0** | **0** |
| 25 | GEE with robust Wald test | **0.00001** | **0** | **0** |
| 100 | GEE with GST | **0.00009** | **0** | **0** |
| 100 | GEE with Wald test | **0.00001** | **0** | **0** |
| - | Independent GEE with Wald test | 0.012 | 0.0011 | 0.00010 |
| - | Wilcoxon rank-sum test | **0.017** | **0.0016** | **0.00017** |
| - | z-test | **0.017** | **0.0021** | **0.00020** |

**Table 2.** Runtime comparison in real ST datasets and simulated ST data.

| Methods | Real Data |  |  |  | Simulated Data |  |  |  |
| --- | --- | --- | --- | --- | --- | --- | --- | --- |
|  | Breast Cancer<br>(3,447 spots, 3,000 genes) |  | Prostate Cancer<br>(4,371 spots, 3,000 genes) |  | Mimicking ST data<br>(3,447 spots, 3,000 replications) |  | Double spots<br>(6,898 spots, 3,000 replications) |  |
|  | Run Time | Memory | Run Time | Memory | Run Time | Memory | Run Time | Memory |
| GEE with GST | 200 s | 3.13 GB | 168 s | 2.69 GB | 171 s | 3.76 GB | 217 s | 3.73 GB |
| GEE with robust Wald test | 15 s | 2.25 GB | 9 s | 3.00 GB | 16 s | 2.25 GB | 45 s | 4.12 GB |
| Independent GEE | 15 s | 3.75 GB | 16 s | 3.00 GB | 16 s | 2.25 GB | 23 s | 4.40 GB |
| Wilcoxon rank-sum test | 4 s | - | 8 s | - | 7 s | - | 10 s | - |
| z-test | 9 s | - | 7 s | - | 8 s | - | 15 s | - |

**Figure 2.**
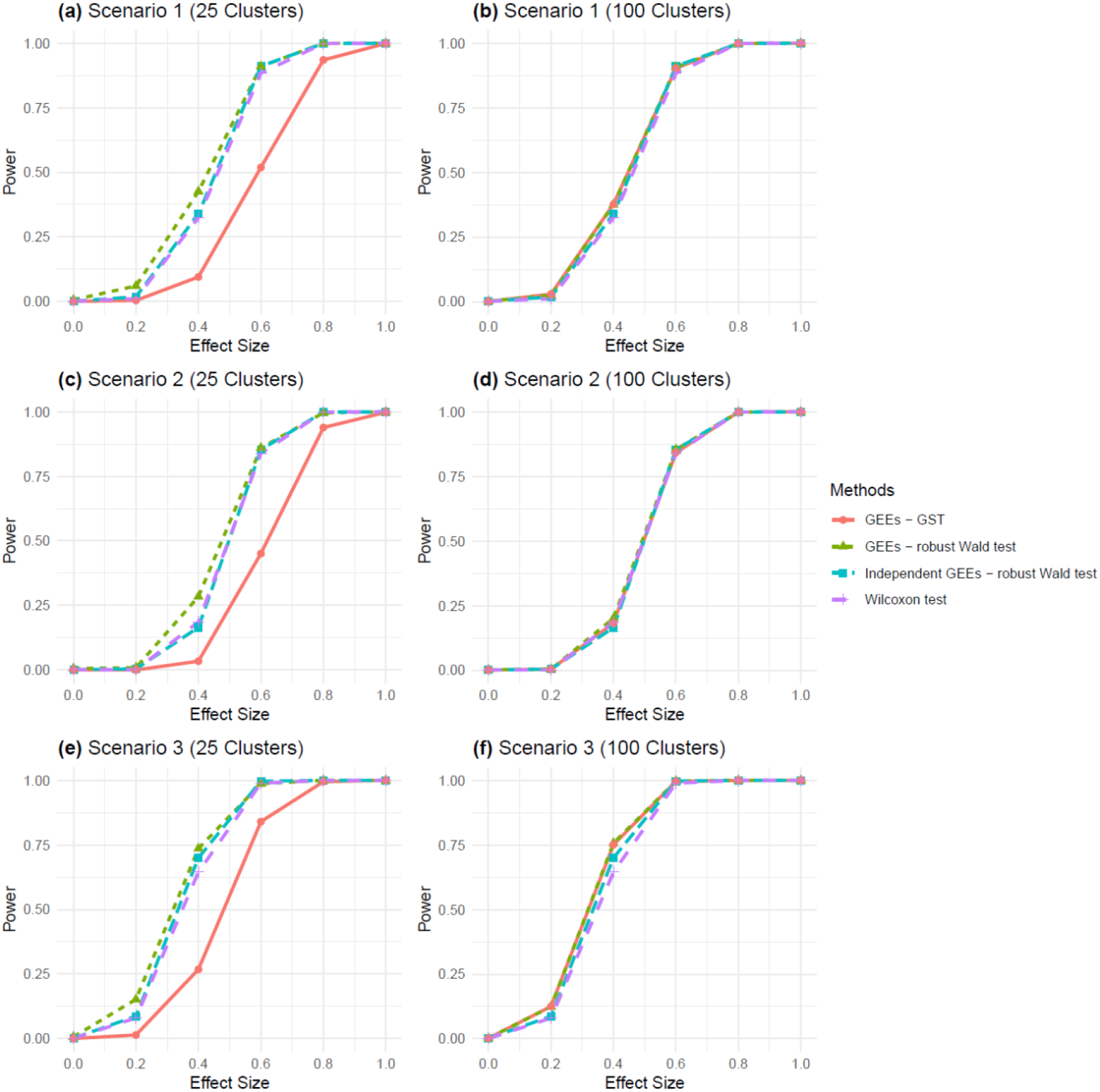
Power comparison of GEE with robust Wald test, GEE with GST, Independent GEE with robust Wald test and Wilcoxon test across different spatial scenarios and spatial cluster numbers. Panels **(a)**–**(f)** represent power curves for three spatial scenarios with 25 and 100 clusters: **(a)** Scenario 1 (25 clusters), **(b)** Scenario 1 (100 clusters), **(c)** Scenario 2 (25 clusters), **(d)** Scenario 2 (100 clusters), **(e)** Scenario 3 (25 clusters), and **(f)** Scenario 3 (100 clusters). Scenarios represent varying spatial correlation and spatial variance: Scenario 1 (weak), Scenario 2 (moderate), and Scenario 3 (strong). Type I error and power were evaluated at a significance level of α = 0.001.

### 3.2. Real Data Applications

In addition to simulation studies, we applied the statistical methods under comparison to two ST datasets for breast cancer and prostate cancer to evaluate their performance in real-world scenarios.

#### 3.2.1. Breast Cancer ST dataset

We applied our methods to a real ST dataset of breast cancer sample provided by 10x Genomics Visium spatial platform (https://www.10xgenomics.com/datasets/human-breast-cancer-block-a-section-1-1-standard-1-1-0). The ST dataset comprises 3,798 spots and 24,923 genes. After retaining the 3,000 most variable genes, the percentage of zeros has a first quartile of 68.0%, a median of 95.5%, and a third quartile of 99.6%. The haematoxylin and eosin (H&E)-stained tissue image with pathology labels is shown in Figure 1a. The analysis used 100 spatial clusters generated via K-means clustering, as illustrated in Figure 1c. Here, the analysis was aimed at identifying the DE genes across pathological grades, specifically comparing ductal carcinoma in situ (DCIS) with fibrous tissue (FT) and invasive carcinoma (IC) with FT. By focusing on breast cancer, we aimed to explore the consistency of our simulation findings using a well-characterized dataset, therefore providing a comprehensive evaluation of the statistical methods under comparison.

The raw dataset was then pre-processed for the analysis by quality control and selection of highly variable genes for downstream analyses. Quality control included the filtration of spots based on low total gene count, resulting in 3,447 spots, and the removal of genes expressed in very few spots to ensure a robust analysis. Using the Seurat software suite, we selected 3,000 most variable genes. This has enabled us to focus on the most informative features of the data and therefore enhance the statistical power in subsequent analyses.

We first generated QQ plots of the genome-wide DE scan p-values (Figure 3) to provide a global assessment of the statistical methods. QQ plots are effective in illustrating discrepancies between observed and expected distributions, particularly in assessing how well each method controls false positives. Among these methods, the Wilcoxon rank-sum test showed the most deviation from the identical line, which might indicate poor calibration of the p-values and possible inflated false positive rates based on findings in the simulation studies (at α = 0.0001). This inflation suggests that users need to interpret the Wilcoxon rank-sum test results with caution when identifying DE genes in ST data where the spatial correlations are pervasive. Excluding the Wilcoxon test, all other tests’ QQ plots behaved similarly (Supplementary Figure S1). In addition, as shown in Supplementary Figures S2 and S3, stratified QQ plots and histograms of p-values by genes’ percentages of zeros for the IC vs FT comparison revealed that the distorted p-value distribution in the overall QQ plot (Figure 3 k) was mainly due to genes with non-sparse counts (percentages of zeros between 0 and 25%, as well as between 25% and 50%), for which the spatial correlations could have a much bigger impact on the calibration of Wilcoxon rank-sum test’s p-values. Of note, the Wilcoxon rank-sum test was shown in the statistical literature to have severely inflated Type I error rates for non-sparse count data under strong correlations (Datta and Satten, 2005).

**Figure 3.**
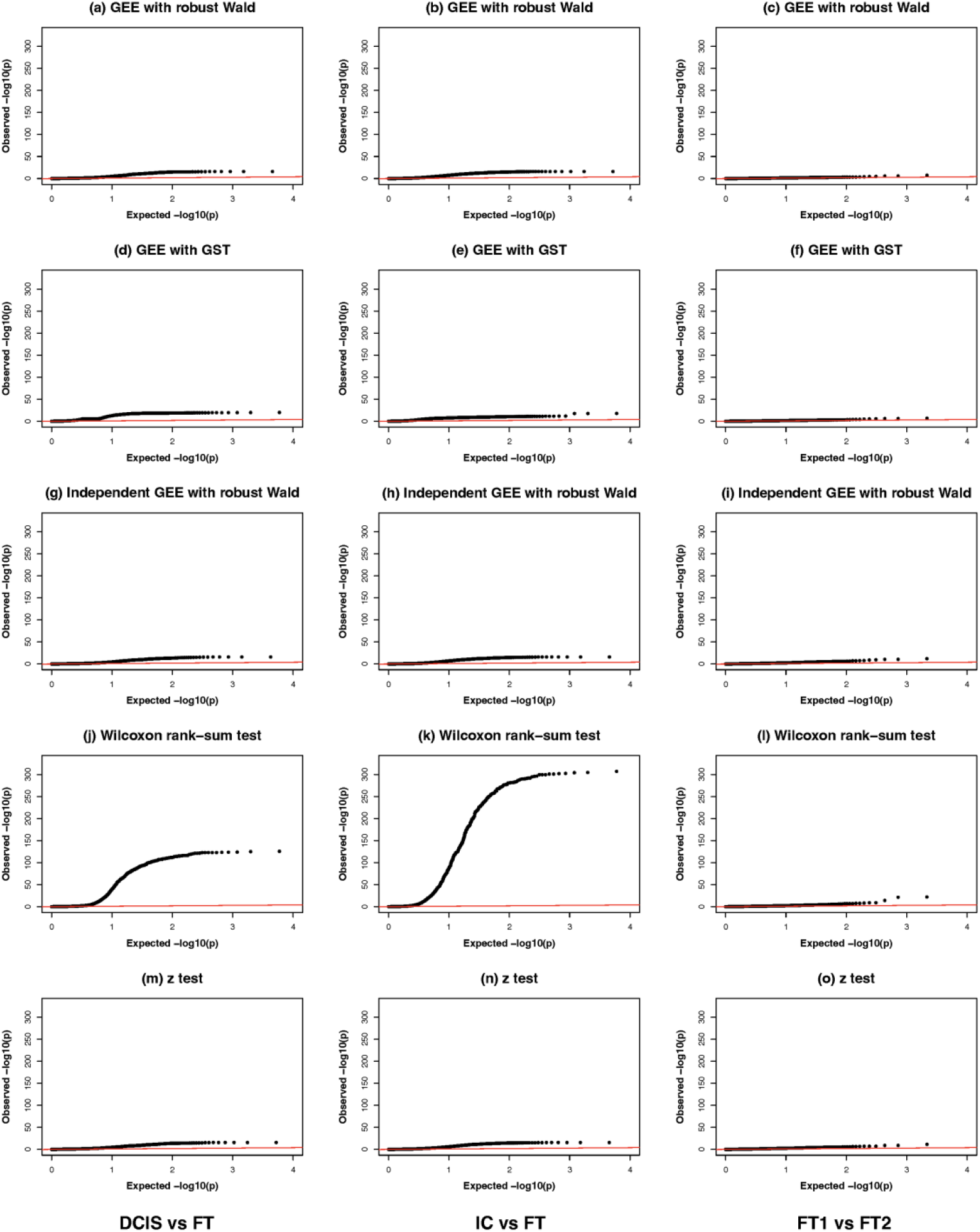
QQ plots of GEE with robust Wald test, GEE with GST, Independent GEE with robust Wald test, Wilcoxon rank-sum test and z-test for breast cancer comparisons: tumor-controls (DCIS vs FT: **(a), (d), (g), (j), (m)**; IC vs FT: **(b), (e), (h), (k), (n)**) and control-control (FT1 vs FT2: **(c), (f), (i), (l), (o)**). The red line represents the expected distribution under the null hypothesis. DCIS: ductal carcinoma in situ; IC: invasive carcinoma; FT: fibrous tissue.

Given that the ground truth of differential expression status of genes is unknown in real datasets, we adopted an internal negative control strategy by comparing the control versus control samples to identify possible false positive DE genes. This approach allows us to indirectly estimate the precision for each method individually without any predefined ground truth. Specifically, as shown in Supplementary Figure S4, we divided the fibrous tissue in the upper right corner of the slide into two subsets (FT1 and FT2) and compared the two subsets against each other. Of note, there has been extensive literature on cancer “field effect”, i.e., molecular and cellular alterations that occur in histologically normal-appearing tissue surrounding a tumor, predisposing that area to cancer development or recurrence (Bondaruk et al., 2022; Guo et al., 2024; Lee et al., 2025). As both FT subsets neighbored with IC and DCIS tissues, we expected that possible differential field effects would be minimal between FT1 and FT2. As shown in the QQ plots for the control-control comparison (Figure 3 (c), (f), (i), (l), and (o)), all the tests behaved similarly, largely following the identical line. As further detailed in Supplementary Figure S5, the Wilcoxon rank-sum test deviated from the identical line the most compared to the other tests. We further used the Venn diagrams to visualize the overlapping significant genes after the Bonferroni correction for FWER = 0.05 between Wilcoxon, GEE with GST and Independent GEE. As shown in Supplementary Figure S6, both Wilcoxon test and Independent GEE identified dozens of significant genes with substantial overlapping, while the GEE with GST’s significant genes were a subset of the Independent GEE’s, consistent with the observations in QQ plots of the real data analysis and the simulation study that the former tended to be more conservative than the latter.

We next conducted biological pathway enrichment analysis on the gene sets identified as differentially expressed by the Wilcoxon rank-sum test, GEE GST and Independent GEE test. For each test, we obtained the significant DE genes by the Bonferroni correction at FWER = 0.05 and retained the top 500 DE genes for pathway enrichment analysis if there were more than 500. The enriched Kyoto Encyclopedia of Genes and Genomes (KEGG) pathways using g:Profiler (Kolberg et al., 2023) at FDR = 0.05 are presented in Figure 4 a, b and c using dot plots.

**Figure 4.**
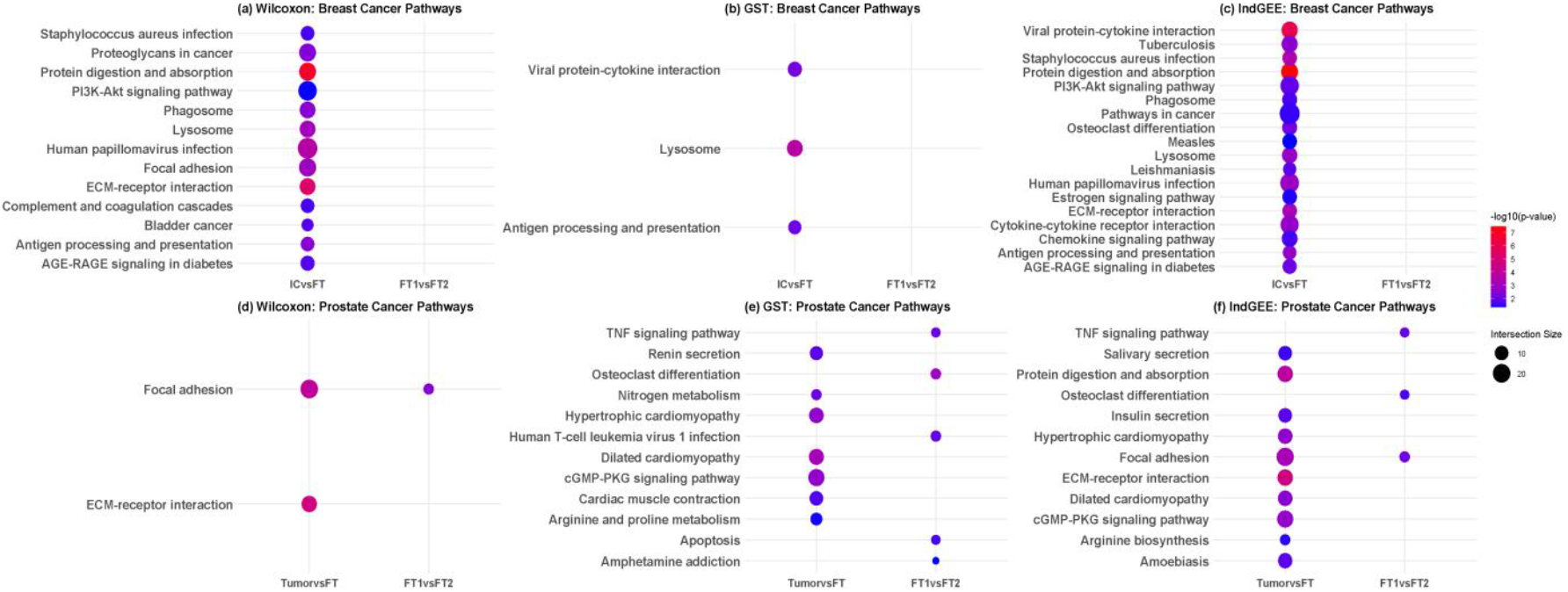
Dot plots of enriched KEGG pathways identified in breast and prostate cancer. Pathways identified by (a) the Wilcoxon rank-sum test, (b) GEE with GST test, and (c) Independent GEE test in the breast cancer dataset, and by the three tests ((d), (e), (f)) in the prostate cancer dataset. IC: invasive carcinoma; FT: fibrous tissue; FT1 vs FT2 (control-vs-control) comparison.

For the IC vs FT comparison, across Wilcoxon rank-sum test, GEE with GST, and Independent GEE, breast cancer exhibits a recurrent signature combining immune/infection-related processes, core oncogenic signaling, and extracellular matrix (ECM) remodeling. Wilcoxon test highlights PI3K–Akt signaling (Vivanco & Sawyers, 2002), focal adhesion (Jin & Varner, 2004), ECM–receptor interaction (Lu et al., 2011), and proteoglycans in cancer (Iozzo & Sanderson, 2011), alongside immune/infectious pathways (e.g., Staphylococcus aureus infection, human papillomavirus infection, phagosome, lysosome, complement/coagulation). GEE with GST narrows to viral protein–cytokine interaction, lysosome, and antigen processing/presentation, indicating targeted enrichment in antigen handling and host–pathogen signaling. Independent GEE identifies the broadest program, extending to estrogen signaling, cytokine–cytokine receptor and chemokine signaling, and multiple infection-related terms, collectively pointing to immune regulation, hormonal control, and matrix–adhesion dynamics as coexisting axes in breast tumor biology. On the other hand, no significant pathway was identified for the FT1 vs FT2 comparison by any of the three tests, consistent with the QQ plots (Supplementary Figure S5) and the expectation of no major biological difference between FT1 and FT2.

#### 3.2.2. Prostate Cancer ST dataset

We extended our comparison further by applying the statistical methods to a prostate cancer ST data, publicly available from 10x Genomics Visium spatial platform (https://www.10xgenomics.com/datasets/human-prostate-cancer-adenocarcinoma-with-invasive-carcinoma-ffpe-1-standard-1-3-0). The ST dataset comprises 4,371 spots and 16,907 genes. After retaining the 3,000 most variable genes, the percentage of zeros has a first quartile of 50.0%, a median of 80.4%, and a third quartile of 97.4%. Figure 1b shows the H&E-stained tissue image with pathology labels. Figure 1d illustrates the 100 spatial clusters generated via K-means clustering. It was of interest to identify DE genes between the tissue types, comparing invasive carcinoma (IC) against adjacent non-cancerous fibro-muscular tissue (FT). The raw data was preprocessed, including quality control and the selection of 3,000 most variable genes using the Seurat pipeline.

As in the breast cancer application, we first created QQ plots of genome-wide DE scans to compare each statistical method’s performance in controlling false positives. As shown in Supplementary Figure S7, the QQ plots showed poor calibration of the p-values for the Wilcoxon rank-sum test. Furthermore, as shown in Supplementary Figures S8 and S9, stratified QQ plots and histograms of p-values confirmed that the distorted p-value distribution was mainly driven by genes with non-sparse counts (percentages of zeros between 0 and 25%, as well as between 25% and 50%), corroborating with the findings in the breast cancer analysis and the simulation studies. This further supports our cautionary note on the Wilcoxon rank-sum test for spatial transcriptomic data analysis where stringent control of false positives is essential.

We then resorted to a similar internal negative control strategy as in the analysis of the breast cancer dataset. As shown in Supplementary Figure S4 (b), the prostate cancer sample was more heterogeneous with intertwining tissues of different pathological grades/classifications compared with the breast cancer sample. So we expected some biological difference at the gene expression/molecular level between FT1 and FT2 in the prostate cancer dataset. QQ plots showed that the observed -log10(p-value) deviated more from the identical line than those in the breast cancer dataset, with the Wilcoxon rank-sum test the most deviating and the GEE with GST the least deviating (Supplementary Figure S10). We further used the Venn diagrams to visualize the overlapping significant genes after the Bonferroni correction for FWER = 0.05 between Wilcoxon, GEE with GST and Independent GEE. As shown in Supplementary Figure S11, similar to the breast cancer dataset, both Wilcoxon and Independent GEE identified dozens of significant genes with substantial overlapping, while the GEE with GST’s significant genes were a subset of the Independent GEE’s, consistent with the observations in QQ plots of the real data analysis and the simulation study that the former tended to be more conservative than the latter. Furthermore, all three tests identified focal adhesion as a significant enriched pathway, which was also identified in the IC vs FT comparison (Figure 4 d, e and f). In addition, GEE with GST and Independent GEE identified the TNF signaling pathway, suggesting possible differential proinflammatory/immune response between FT1 and FT2 due to the field effect.

### 3.3. Comparison between simulated data and real data

As shown in Supplementary Figure S12, while both real and simulated ST datasets exhibit substantial sparsity, the degree and distribution differ across spots and genes. In the real data, the 3,000 most variable genes showed varying zero percentages and spatial correlations. In the simulated data, we simulated a single gene across 100,000 replications for a given set of spatial correlation parameters to evaluate the Type I error rates and introducing high-percentage of zeros through a small intercept (*β*_0_) in the GLMM-based data generating model.

Regarding per-gene sparsity, at the gene level, the Visium ST breast cancer dataset displayed a wider range of sparsity, with most genes containing 60–95% zeros and a modest proportion with only 0% to 50% zeros. In the simulated dataset, however, sparsity was more extreme and concentrated, with most replications containing over 90% zeros, reflecting stronger zero inflation by design.

As for per-spot sparsity, in the Visium ST breast cancer dataset, most spots contained approximately 70–90% zeros, with a peak around 80–85%. In contrast, the simulated zero-inflated dataset showed even higher sparsity, with the majority of spots containing over 90% zeros and a right-skewed distribution concentrated near 100%.

Overall, the simulated data successfully reproduced the high sparsity characteristic of real ST data, though with slightly more pronounced zero inflation at both the spot and gene levels. As a result, the real data and simulated data complemented each other, together providing more comprehensive insights into the comparative performance of the various statistical tests.

### 3.4. Computational runtime comparison

We compared the computational efficiency of different methods using both real ST datasets (each containing 3,000 genes) and simulated datasets (each containing 3,000 replications of a single gene from the moderate spatial correlation scenario in Section 3.1). All analyses were performed using 20 CPU cores on an Intel(R) Xeon(R) Gold 6254 processor (3.10 GHz). Across datasets, the GEE with GST required the longest runtime and the largest memory footprint, reflecting the additional computation involved in estimating spatial covariance parameters and matrix inversion (approximately 170 - 220 s and 2.7 - 3.8 GB of memory). The GEE with the robust Wald test and the Independent GEE were substantially faster, completing within 9 - 45 s while using 2 - 4 GB of memory. In contrast, the non-spatial tests, the Wilcoxon rank-sum test and the two-sample z-test, were the most computationally efficient, finishing within 4 - 15 s with negligible additional memory requirements. When the number of spatial spots doubled from 3,447 to 6,898 in the simulated data, runtimes approximately doubled for all methods, but the relative ranking of computational cost remained consistent.

## 4. Discussion

We have performed a comprehensive comparison of statistical methods for identified DE genes in ST data with evaluation of Type I error control, power, and identification of biologically relevant pathways. Through extensive simulations and applications to breast cancer and prostate cancer ST datasets, we compared the robustness of each method in handling the spatial correlations inherent in ST data. We summarize our main findings and recommendations in Table 3. The most commonly used Wilcoxon rank-sum test offers fast computation and is recommended for sparse datasets, though caution is warranted under non-sparse data and high spatial correlation. The two-sample z-test fails to account for spatial dependencies and is not recommended. GEE models vary in performance: while GEE with robust Wald tests may inflate Type I error and GEE with GST may deflate it, Independent GEE provides accurate error control and complements the Wilcoxon rank-sum test. GLMMs, despite modeling spatial correlation via random effects, suffer from convergence issues and are computationally intensive, limiting their utility. Overall, Independent GEE and Wilcoxon rank-sum tests emerge as the most reliable choices for ST count data analysis.

**Table 3.** Comparative performance of various statistical tests for detecting DE genes in ST data.

| Method | Model Type | Accounts for Correlation? | Type I Error Control | Computational runtime | Software | Recommended for ST count data |
| --- | --- | --- | --- | --- | --- | --- |
| Wilcoxon Rank-Sum Test | Nonparametric rank-based | No | Could inflate or deflate Type I error rate depending on sparsity and spatial correlation patterns of the count data | Very fast | Seurat; SpatialGE; SpatialGEE | Yes, but caution with non-sparse data and/or high spatial correlations |
| Z-Test (Two-Sample) | Large-sample parametric | No | Could inflate or deflate Type I error rate depending on spatial correlation patterns of the count data | Very fast | “z.test” function in R package “BSDA” | No |
| GEE with GST | Generalized Estimating Equations | Yes - incorporates spatial correlation via sandwich variance estimator | Depends on the number of pre-specified clusters $m$; tends to deflate the Type I error rate, and, thus, reduced power | Slow | SpatialGEE | No |
| GEE with Robust Wald Test | Generalized Estimating Equations | Yes – incorporates spatial correlation via sandwich variance estimator | Depends on the number of pre-specified clusters $m$; tends to inflate the Type I error rate | Fast | SpatialGEE | No |
| Independent GEE | Generalized Estimating Equations | Yes – incorporates spatial correlation via sandwich variance estimator | Accurate control | Fast | SpatialGEE | Yes, complementary to Wilcoxon rank-sum test |
| GLMM | Generalized linear mixed model | Yes – incorporate spatial correlation via spatial random effects | Not evaluable due to convergence issue | Very slow; convergence issue | R package “spaMM” | No |

The GEE model uses the working independence correlation for spots within each cluster; it then uses the empirical covariances within and between clusters to correct for the likely incorrect working independence assumptions within and between clusters in the “sandwich” robust covariance estimator framework that remains valid asymptotically as long as the number of clusters *m* is large. We have shown that (1) *m* needs to be at least 100 to control the Type I error rate in the GEE GST and GEE robust Wald test (Section 3.1 and 3.2), and (2) independent GEE with the robust standard error, i.e., each spot forms its own cluster without the need for K-means pre-clustering, achieved superior Type I error control and computational efficiency. We therefore recommend the independent GEE method to be a useful complement to the Wilcoxon rank-sum test in identifying DE genes in ST count data.

Several ST platforms are available and generate data with different characteristics, such as 10x Visium ST, 10x Xenium, 10x Visium HD, Slide-Seq, MerFISH, and CosMX. Our proposed GST approach is in principle widely applicable to all these platforms, similar to the Wilcoxon rank-sum test implemented in Seurat. We specifically used two 10x Visium ST datasets in our real data applications, as 10x Visium is thus far the most widely adopted platform. Future research is warranted to study the robustness of Independent GEE and Wilcoxon rank-sum test across other ST technologies.

## Data and Code Availability

We have implemented our proposed methods in R package ‘SpatialGEE’ available at https://github.com/yishan03/SpatialGEE. The spatial transcriptomics datasets of breast cancer and prostate cancer analyzed here are available on 10x Genomics websites: https://www.10xgenomics.com/datasets/human-breast-cancer-block-a-section-1-1-standard-1-1-0 and https://www.10xgenomics.com/datasets/human-prostate-cancer-adenocarcinoma-with-invasive-carcinoma-ffpe-1-standard-1-3-0.

## Supporting Information

The supplementary materials include Supplementary Figures S1 to S12.

## Acknowledgments

This work was supported by Cancer Prevention and Research Institute of Texas (CPRIT) grant RP230166 (to PW). PW was partially supported by National Institutes of Health (NIH) grants P50CA217674 and P01CA296429. The authors declare no conflict of interest.

