## Supplementary Figures S1 to S12 for "A comparative study of statistical methods for identifying differentially expressed genes in spatial transcriptomics"

**Figure S1.** QQ plots of (a) GEE with robust Wald test, (b) GEE with GST, (c) Independent GEE with robust Wald test, and (d) two-sample z-test (excluding the Wilcoxon rank-sum test) for IC vs FT comparison in the breast cancer ST dataset.

**Figure S2.** QQ plots of the Wilcoxon rank-sum test p-values stratified by genes' percentages of zeros for the IC vs FT comparison in the breast cancer ST dataset.

**Figure S3.** Histograms of the Wilcoxon rank-sum test and Independent GEE p-values stratified by genes' percentages of zeros for the IC vs FT comparison in the breast cancer ST dataset.

**Figure S4.** FT1 vs FT2 comparison in breast cancer ((a) and (c)) and prostate cancer ((b) and (d)) ST datasets.

**Figure S5.** QQ plots of various tests for FT2 vs FT2 comparison in the breast cancer ST dataset. FT: fibrous tissue.

**Figure S6.** Venn diagram of the significant genes after the Bonferroni correction for FWER = 0.05 between Wilcoxon rank-sum test, GEE with GST and Independent GEE for FT2 vs FT2 comparison in the breast cancer ST dataset.

**Figure S7.** QQ plots of GEE with robust Wald test, GEE with GST, Independent GEE with robust Wald test, Wilcoxon rank-sum test and z-test for prostate cancer comparisons.

**Figure S8.** QQ plots of the Wilcoxon rank-sum test p-values stratified by genes' percentages of zeros for the IC vs FT comparison in the prostate cancer ST dataset.

**Figure S9.** Histograms of the Wilcoxon rank-sum test and Independent GEE p-values stratified by genes' percentages of zeros for the IC vs FT comparison in the prostate cancer ST dataset.

**Figure S10.** QQ plots of various tests for FT2 vs FT2 comparison in the prostate cancer ST dataset.

**Figure S11.** Venn diagram of the significant genes after the Bonferroni correction for FWER = 0.05 between Wilcoxon rank-sum test, GEE with GST and Independent GEE for FT2 vs FT2 comparison in the prostate cancer ST dataset.

**Figure S12.** Comparison between simulated data and breast cancer ST real data regarding (a) per-gene/replication sparsity and (b) per-cell/spot sparsity.

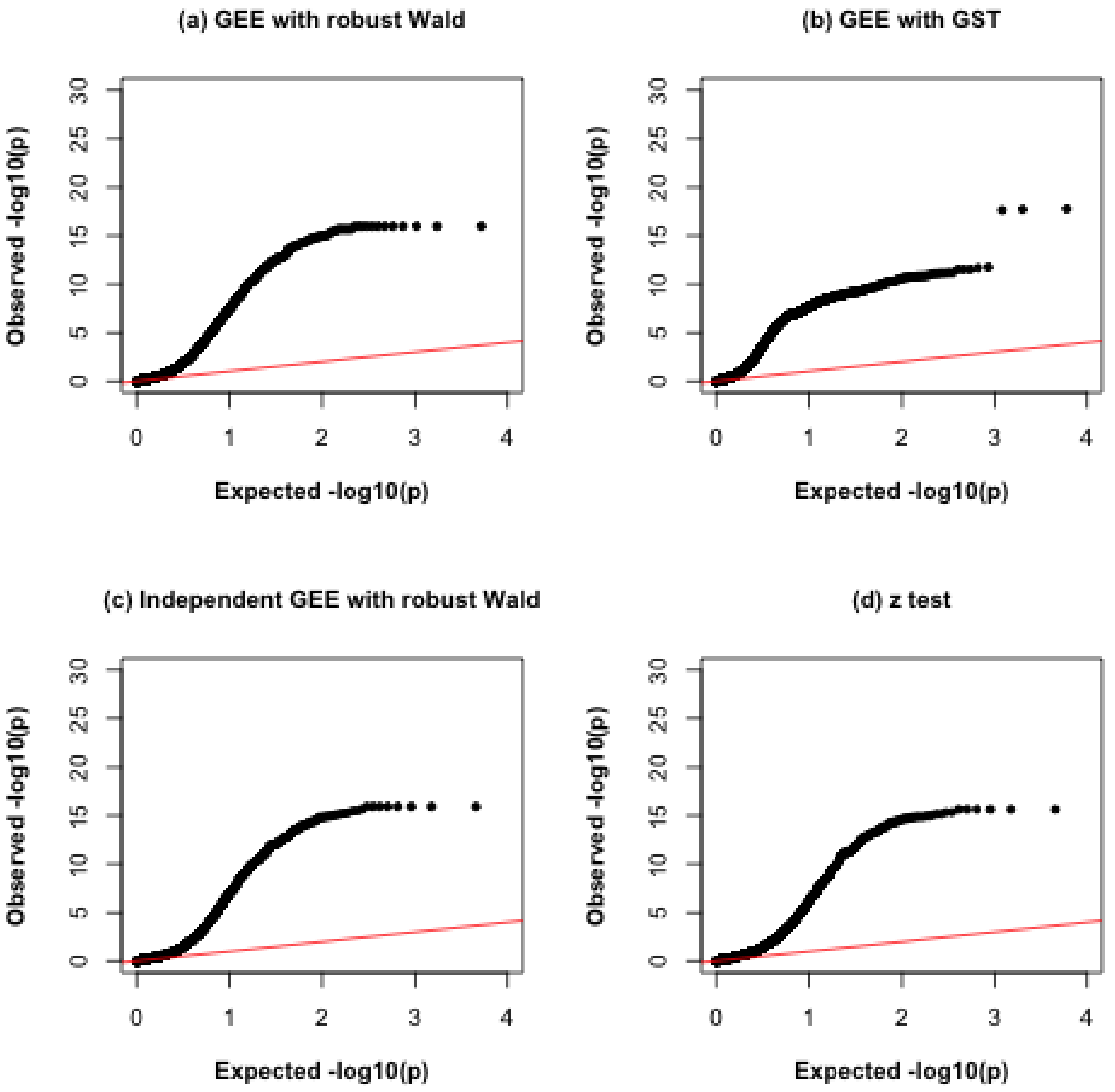


**Figure S1.** QQ plots of (a) GEE with robust Wald test, (b) GEE with GST, (c) Independent GEE with robust Wald test, and (d) two-sample z-test (excluding the Wilcoxon rank-sum test) for IC vs FT comparison in the breast cancer ST dataset. IC: invasive carcinoma; FT: fibrous tissue.

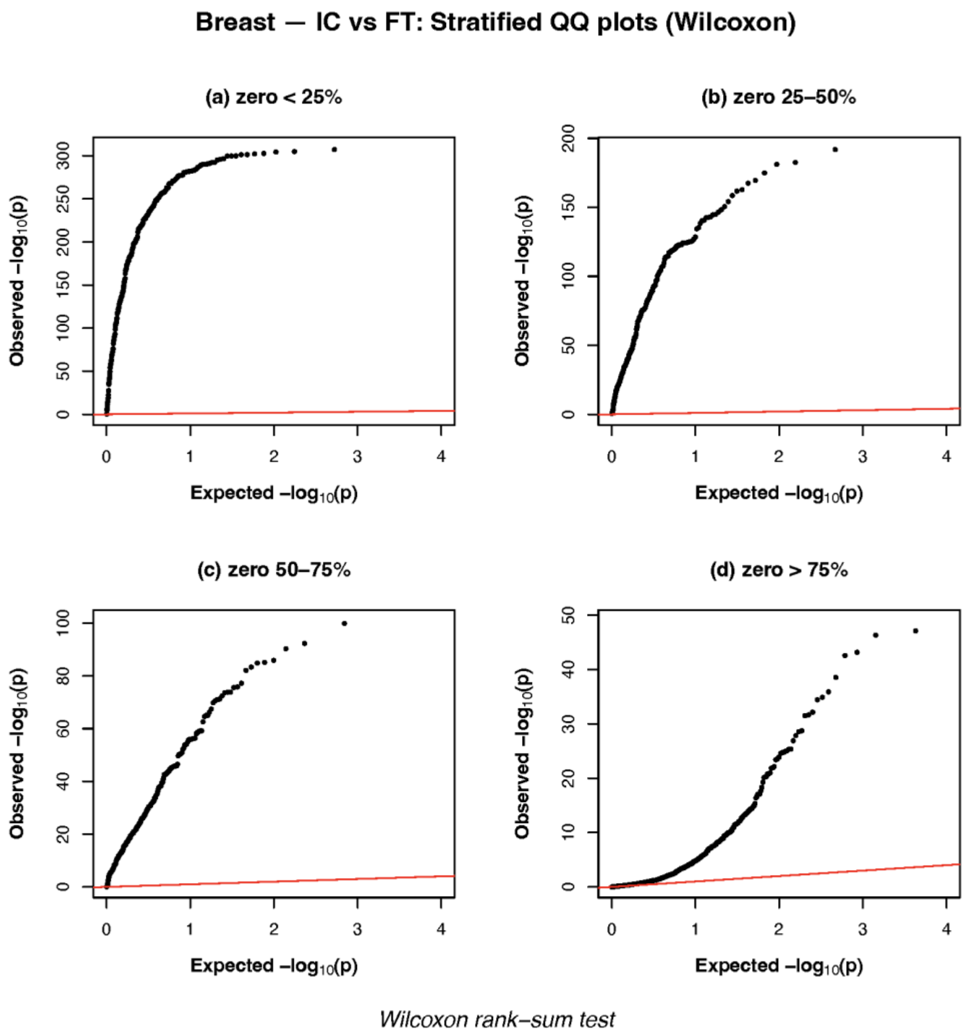


**Figure S2.** QQ plots of the Wilcoxon rank-sum test p-values stratified by genes' percentages of zeros for the IC vs FT comparison in the breast cancer ST dataset. Note that the Y limits in the QQ plots were at 300, 200, 100 and 50, respectively.

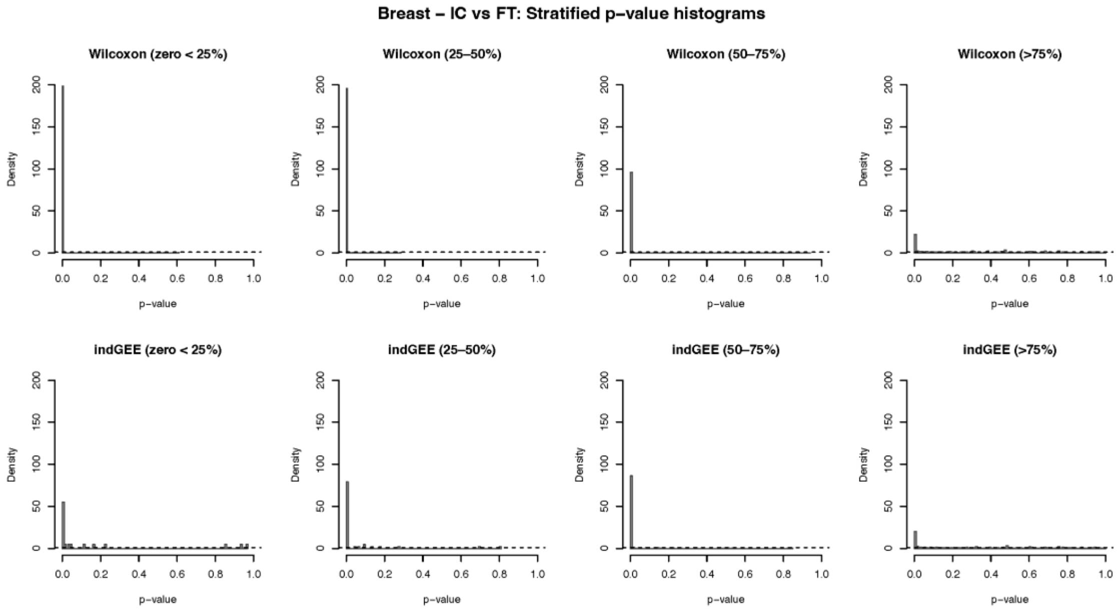


**Figure S3.** Histograms of the Wilcoxon rank-sum test and Independent GEE p-values stratified by genes' percentages of zeros for the IC vs FT comparison in the breast cancer ST dataset.

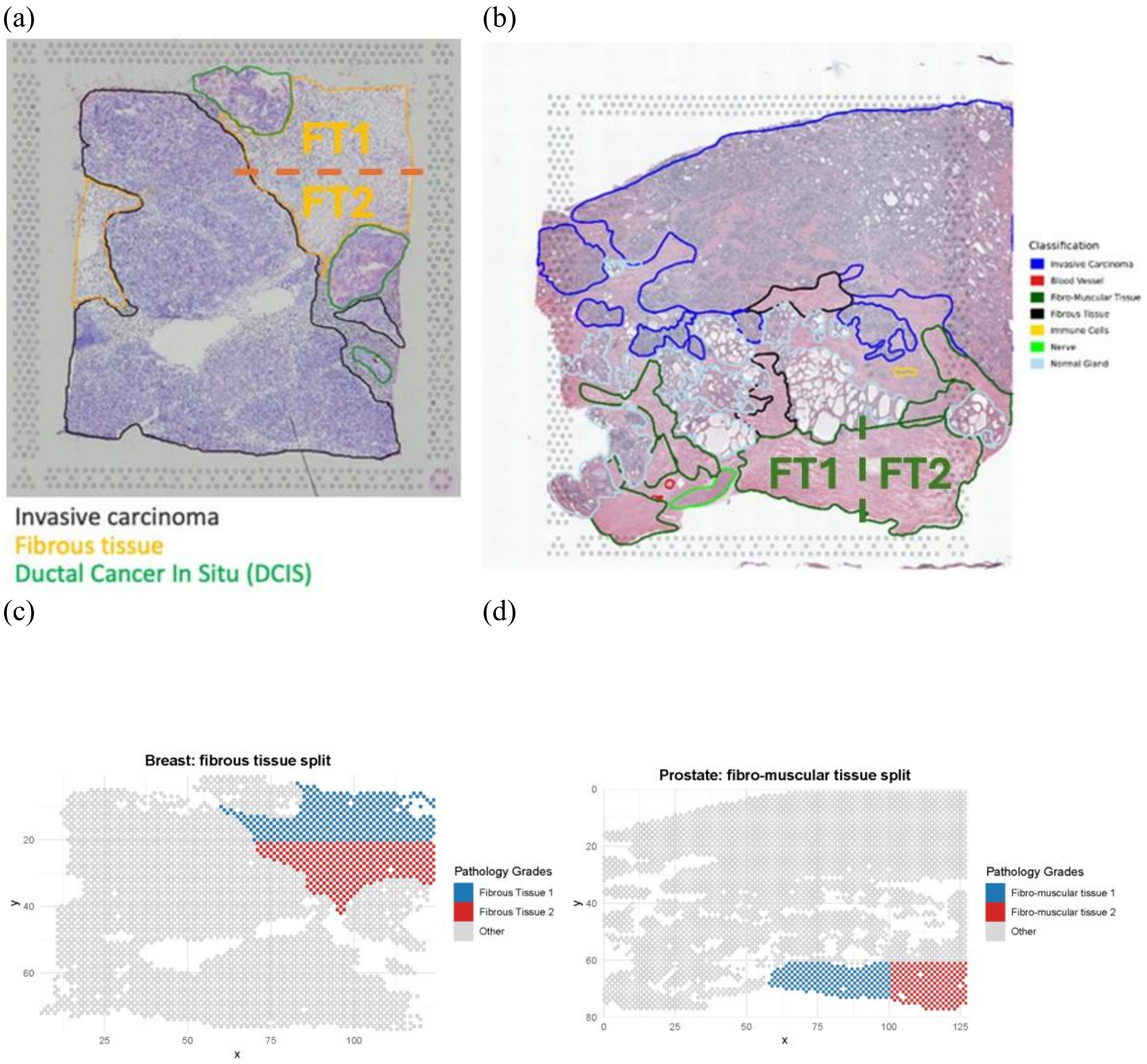


**Figure S4.** FT1 vs FT2 comparison in breast cancer ((a) and (c)) and prostate cancer ((b) and (d)) ST datasets.

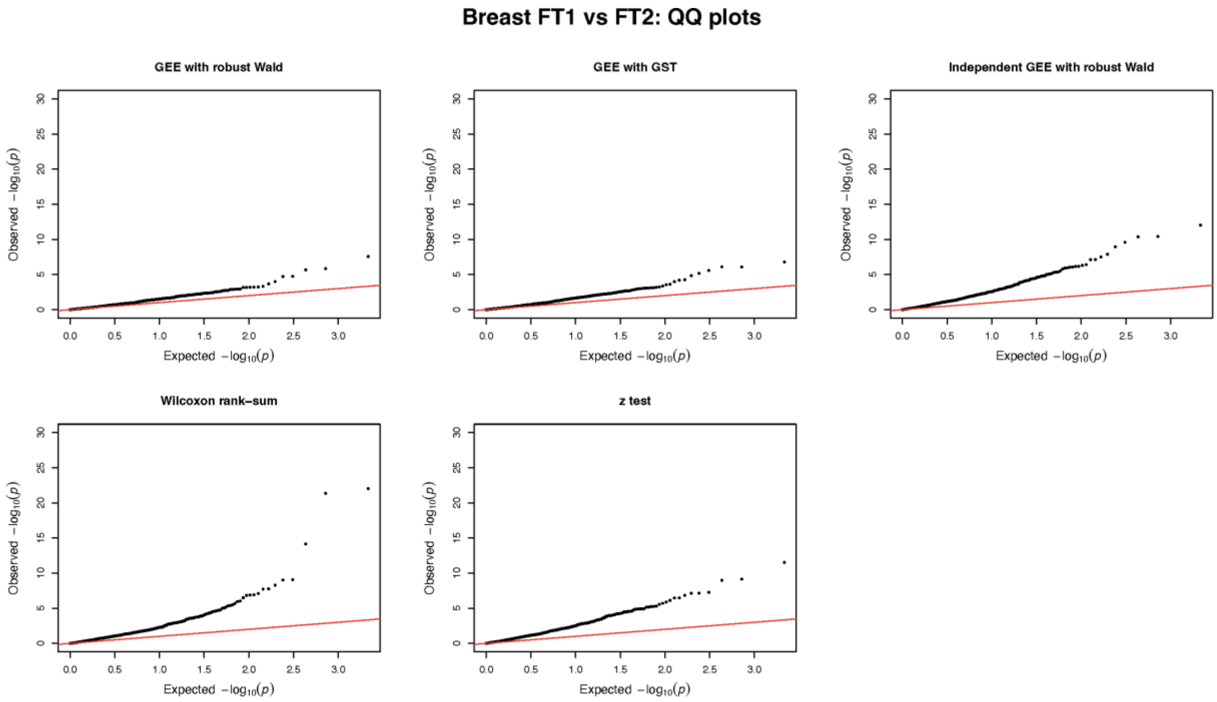


**Figure S5.** QQ plots of various tests for FT2 vs FT2 comparison in the breast cancer ST dataset. FT: fibrous tissue.

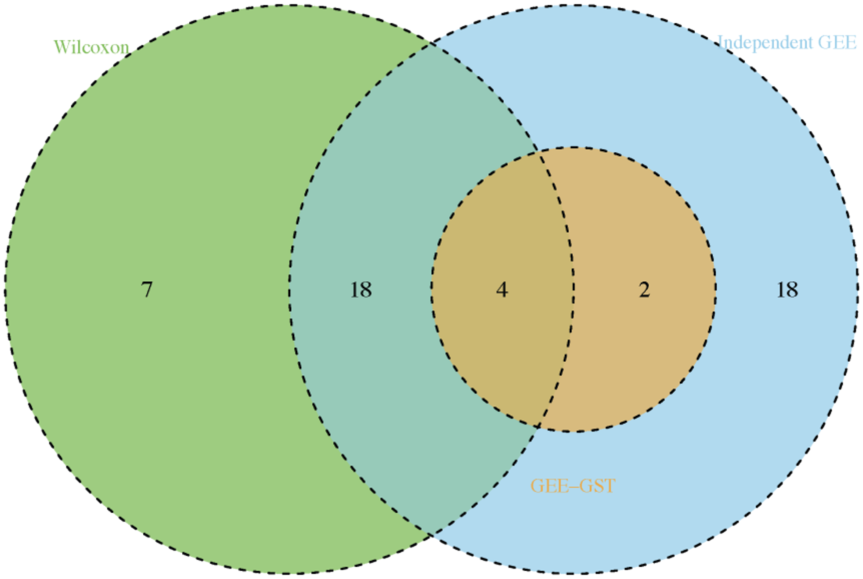


**Figure S6.** Venn diagram of the significant genes after the Bonferroni correction for FWER = 0.05 between Wilcoxon rank-sum test, GEE with GST and Independent GEE for FT2 vs FT2 comparison in the breast cancer ST dataset.

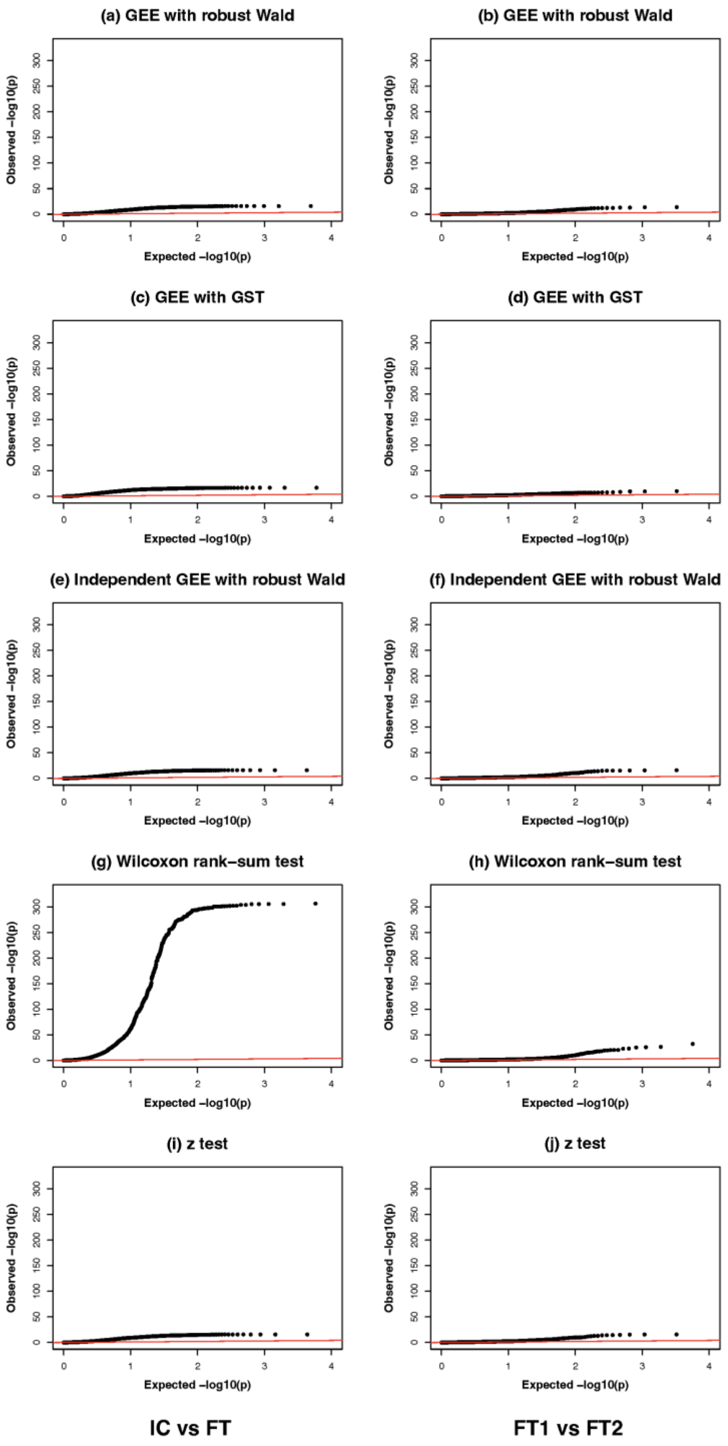


**Figure S7.** QQ plots of GEE with robust Wald test, GEE with GST, Independent GEE with robust Wald test, Wilcoxon rank-sum test and z-test for prostate cancer comparisons: tumor-controls (IC vs FT: **(a)**, **(c)**, **(e)**, **(g), (i)**) and control-control (FT1 vs FT2: **(b)**, **(d)**, **(f)**, **(h), (j)**). The red line represents the expected distribution under the null hypothesis. IC: invasive carcinoma; FT: fibrous tissue.

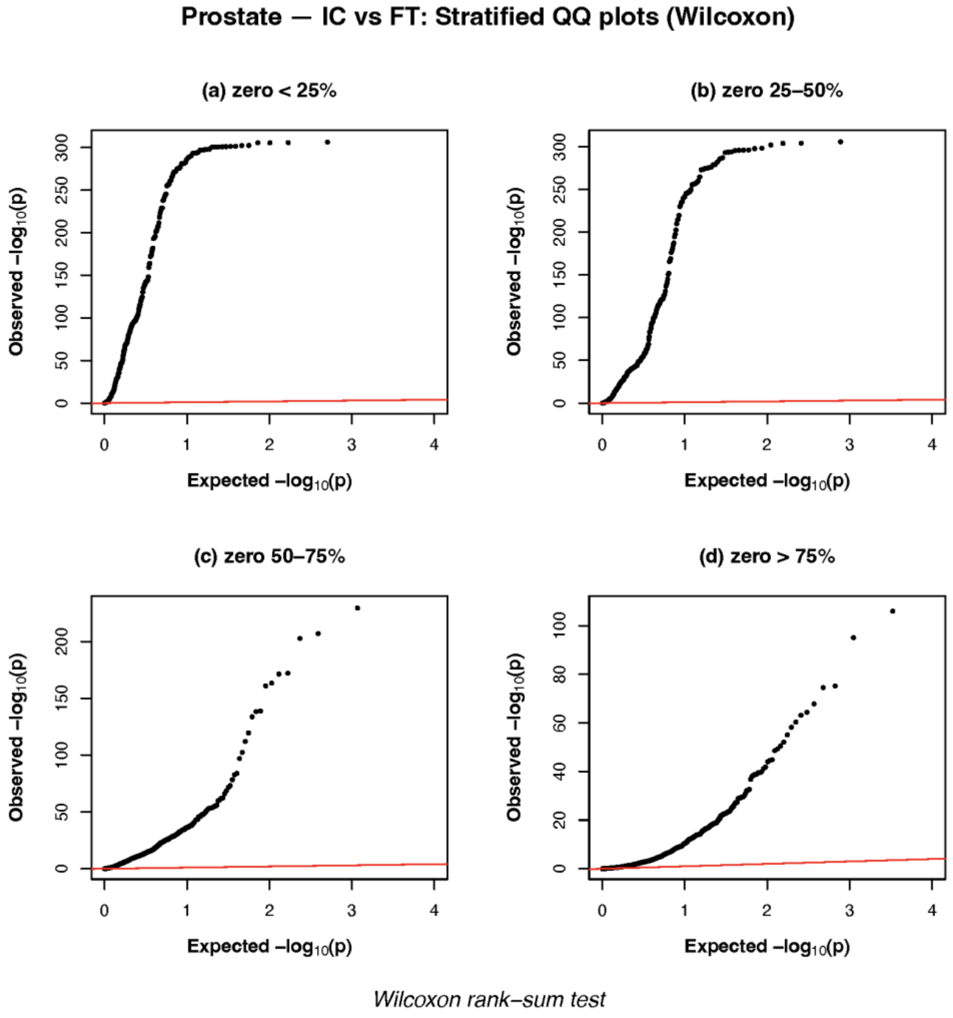


**Figure S8.** QQ plots of the Wilcoxon rank-sum test p-values stratified by genes' percentages of zeros for the IC vs FT comparison in the prostate cancer ST dataset. Note that the Y limits in the QQ plots were at 300, 300, 200 and 100, respectively.

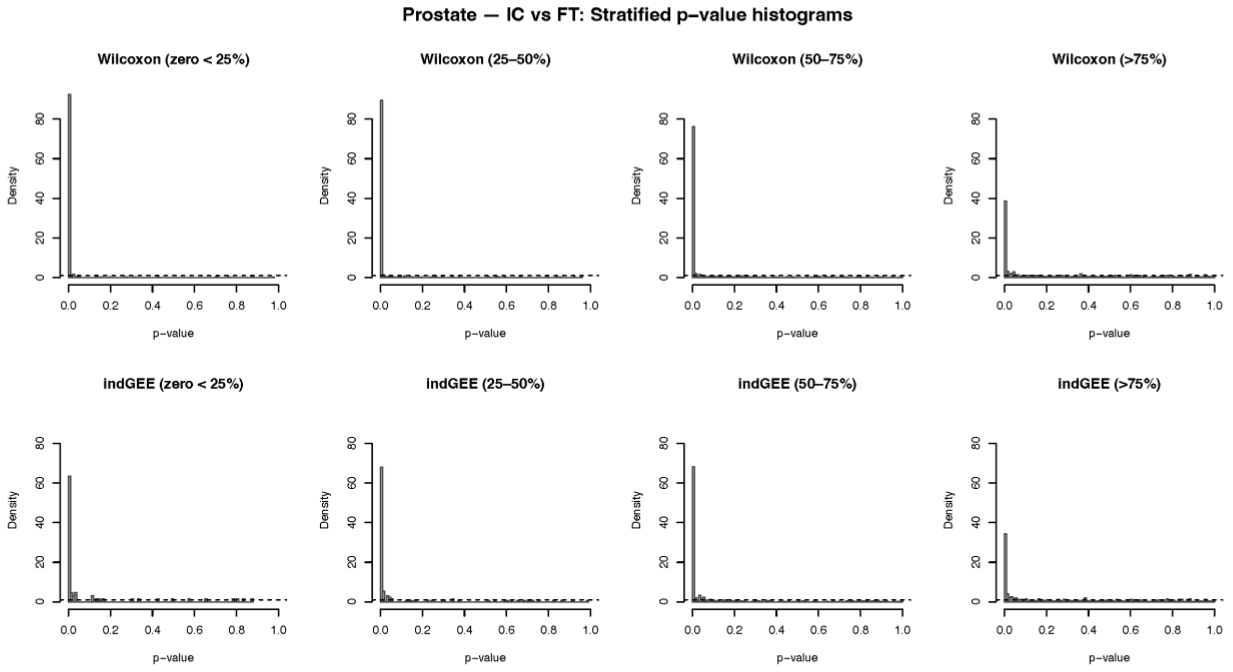


**Figure S9.** Histograms of the Wilcoxon rank-sum test and Independent GEE p-values stratified by genes' percentages of zeros for the IC vs FT comparison in the prostate cancer ST dataset.

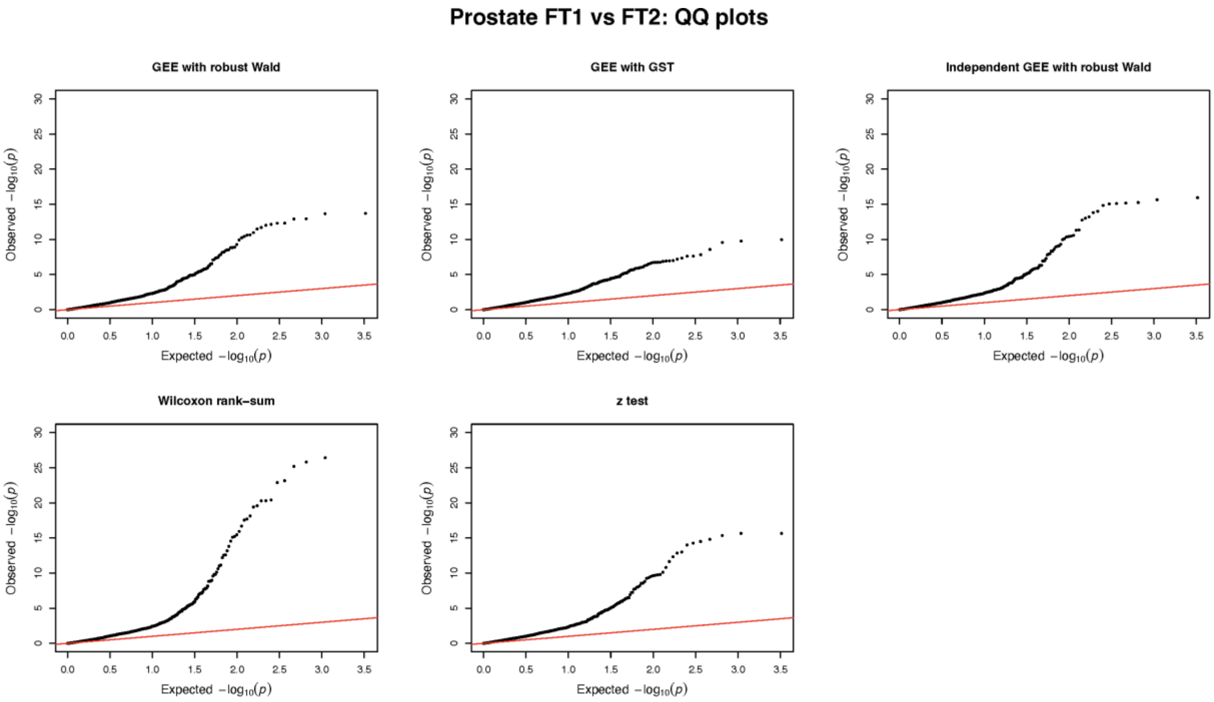


**Figure S10.** QQ plots of various tests for FT2 vs FT2 comparison in the prostate cancer ST dataset. FT: fibrous tissue.

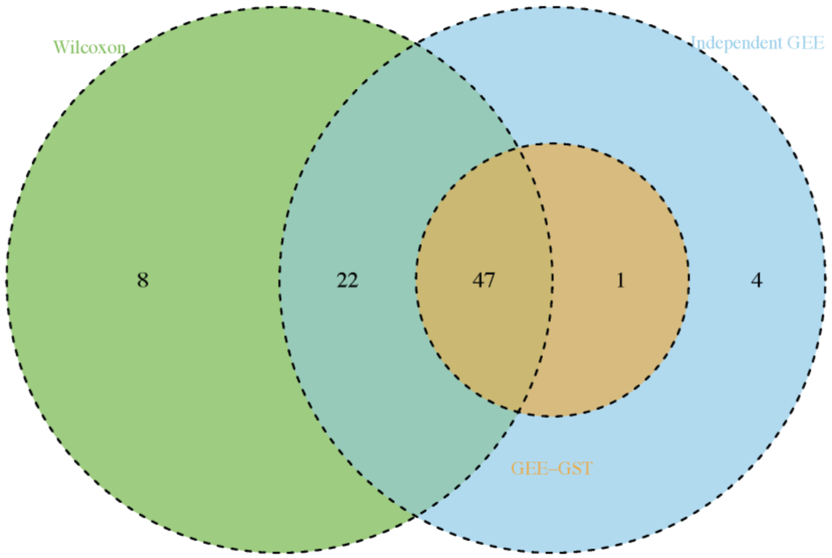


**Figure S11.** Venn diagram of the significant genes after the Bonferroni correction for FWER = 0.05 between Wilcoxon rank-sum test, GEE with GST and Independent GEE for FT2 vs FT2 comparison in the prostate cancer ST dataset.

(a)

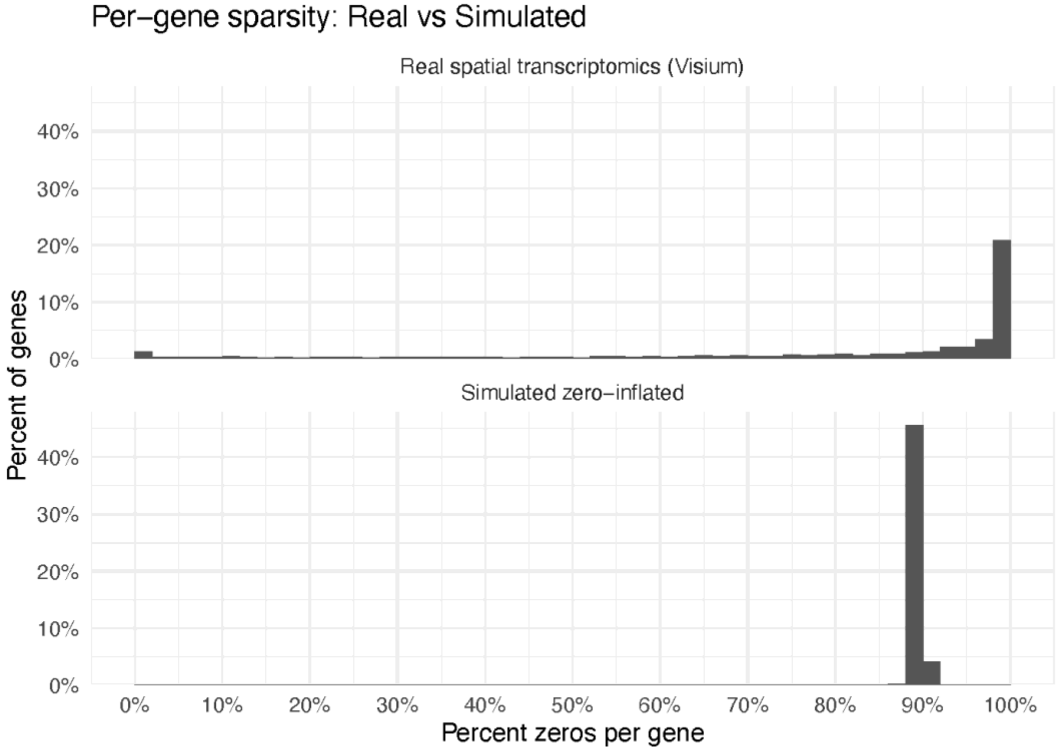


(b)

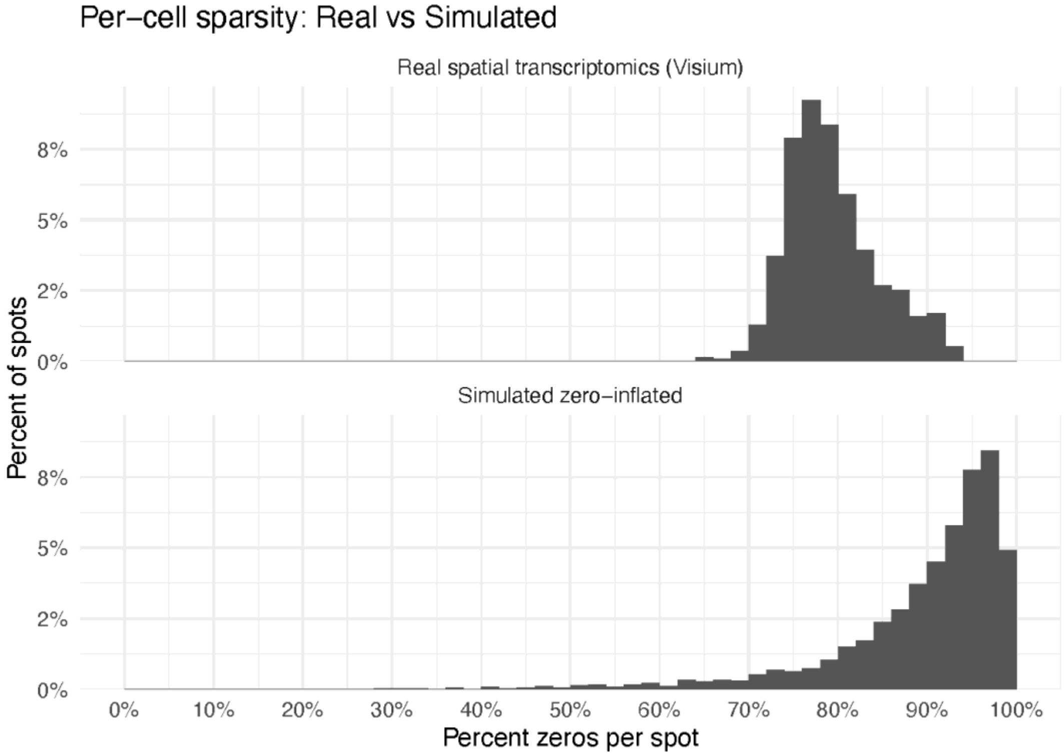


**Figure S12.** Comparison between simulated data and breast cancer ST real data regarding (a) per-gene/replication sparsity and (b) per-cell/spot sparsity.
